# Freely behaving mice can brake and turn during optogenetic stimulation of the Mesencephalic Locomotor Region

**DOI:** 10.1101/2020.11.30.404525

**Authors:** Cornelis Immanuel van der Zouwen, Joël Boutin, Maxime Fougère, Aurélie Flaive, Mélanie Vivancos, Alessandro Santuz, Turgay Akay, Philippe Sarret, Dimitri Ryczko

## Abstract

**Background:** Stimulation of the Mesencephalic Locomotor Region (**MLR**) is increasingly considered as a target to improve locomotor function in Parkinson’s disease, spinal cord injury and stroke. A key function of the MLR is to control the speed of forward symmetrical locomotor movements. However, the ability of freely moving mammals to integrate environmental cues to brake and turn during MLR stimulation is poorly documented.

**Objective/hypothesis:** We investigated whether freely behaving mice could brake or turn based on environmental cues during MLR stimulation.

**Methods:** We stimulated the cuneiform nucleus in mice expressing channelrhodopsin in Vglut2-positive neurons in a Cre-dependent manner (Vglut2-ChR2-EYFP) using optogenetics. We detected locomotor movements using deep learning. We used patch-clamp recordings to validate the functional expression of channelrhodopsin and neuroanatomy to visualize the stimulation sites.

**Results:** Optogenetic stimulation of the MLR evoked locomotion and increasing laser power increased locomotor speed. Gait diagram and limb kinematics were similar during spontaneous and optogenetic-evoked locomotion. Mice could brake and make sharp turns (∼90⁰) when approaching a corner during MLR stimulation in an open-field arena. The speed during the turn was scaled with the speed before the turn, and with the turn angle. In a reporter mouse, many Vglut2-ZsGreen neurons were immunopositive for glutamate in the MLR. Patch-clamp recordings in Vglut2-ChR2-EYFP mice show that blue light evoked short latency spiking in MLR neurons.

**Conclusion:** MLR glutamatergic neurons are a relevant target to improve locomotor activity without impeding the ability to brake and turn when approaching an obstacle, thus ensuring smooth and adaptable navigation.

**Highlights:**

- Mice brake and turn when approaching the arena’s corner during MLR-evoked locomotion
- Speed decrease is scaled to speed before the turn during MLR-evoked locomotion
- Turn angle is scaled to turn speed during MLR-evoked locomotion
- Gait and limb kinematics are similar during spontaneous and MLR-evoked locomotion

## Introduction

Coordination of speed, braking and steering is essential to navigate the environment [1]. In the brainstem, the Mesencephalic Locomotor Region (**MLR**) plays a key role in initiating and controlling locomotion ([2], for review [3]). MLR glutamatergic neurons control locomotor speed from basal vertebrates to mammals (e.g. lamprey [4, 5, 6], salamanders [7], mice [8, 9, 10, 11, 12]). A key function of the MLR is to elicit forward symmetrical locomotion by sending bilateral glutamatergic inputs to reticulospinal neurons that project to the spinal central pattern generator for locomotion (cat [13], lamprey [14, 15], zebrafish [16, 17], salamander [18], mouse [19, 20, 10]). In mammals, MLR commands are relayed by reticulospinal neurons in the lateral paragigantocellular nucleus (**LPGi**) [10].

Steering movements are induced by asymmetrical reticulospinal activity. Increased reticulospinal activity on one side induces ipsilateral steering movements in lamprey [21, 22, 23, 24], zebrafish [25, 26], salamander [27] and rat [28]. In mammals, steering commands are relayed by reticulospinal neurons in the gigantocellular nucleus (**Gi**) that express the molecular marker Chx10. Their unilateral activation evokes ipsilateral braking and turning [29, 30, 31]. These neurons receive no input from the MLR, but a major glutamatergic input from the contralateral superior colliculus (**SC**), a region involved in visuomotor transformations ([32, 33, 34], for review [35]).

The interactions between brainstem substrates controlling speed (MLR-LPGi) and those controlling braking and turning (SC-Gi) are unknown. Whether braking or turning can be done during MLR stimulation is poorly documented in mammals. In mice, MLR-evoked locomotion was recorded on a trackball [8, 9], treadmill [11] or in a linear corridor [12], but the ability to integrate environmental cues that modify the direction of motion was not studied. In decerebrated cats held over a treadmill oriented in various directions, MLR stimulation generated well-coordinated locomotion only when the treadmill was going in front-to-rear direction, suggesting that the MLR could only generate forward motion [36].

Here, we examined whether freely behaving mice can brake or turn by integrating environmental cues during optogenetic stimulation of the MLR. In mice expressing channelrhodopsin in neurons positive for the vesicular glutamatergic transporter 2 (Vglut2-ChR2-EYFP mice), we targeted the cuneiform nucleus (**CnF**), the MLR subregion that controls the largest range of speeds [11, 12]. We used deep learning to detect locomotor movements in a linear corridor and in an open field arena. It is relevant to determine whether MLR-evoked locomotion can be dynamically adapted to the environment, as MLR stimulation is explored to improve locomotor function in Parkinson’s disease [37, 38, 39, 40] and in animal models of spinal cord injury [41, 42] and stroke [43]. We focused on the CnF, which is increasingly considered as the optimal subregion to target within the MLR [44].

## Materials and Methods

The procedures were in accordance with the guidelines of the Canadian Council on Animal Care and approved by the animal care and use committees of the Université de Sherbrooke.

### Animals

Thirteen mice were used. We used Vglut2-Cre mice (Jackson laboratories, #028863, Vglut2-ires-cre knock-in (C57BL/6J)) [45] (Fig. 1A-B), ChR2-EYFP-lox mice (Ai32 mice, Jackson laboratory, #024109, B6.Cg-*Gt(ROSA)26Sor^tm32(CAG-COP4*H134R/EYFP)Hze^*/J) [46] (Fig. 1B), and ZsGreen-lox mice (Ai6 mice, Jackson laboratory, #007906, B6.Cg-*Gt(ROSA)26Sor^tm6(CAG-ZsGreen1)Hze^*/J) [46] (Fig. 1A). We crossed homozygous Vglut2-Cre mice with homozygous ChR2-EYFP-lox mice to obtain the double heterozygous Vglut2-ChR2-EYFP mice. We crossed homozygous Vglut2-Cre mice with homozygous ZsGreen-lox mice to obtain the double heterozygous Vglut2-ZsGreen mice. Animals had *ad libitum* access to food and water, with lights on from 6 AM to 8 PM. Mice were 16-36 weeks old for in vivo optogenetics (3 males, 2 females), 10-18 weeks old for neuroanatomy (1 male, 5 females), and 15-23 days old for patch-clamp experiments (1 male, 1 undetermined).

**Figure 1.**
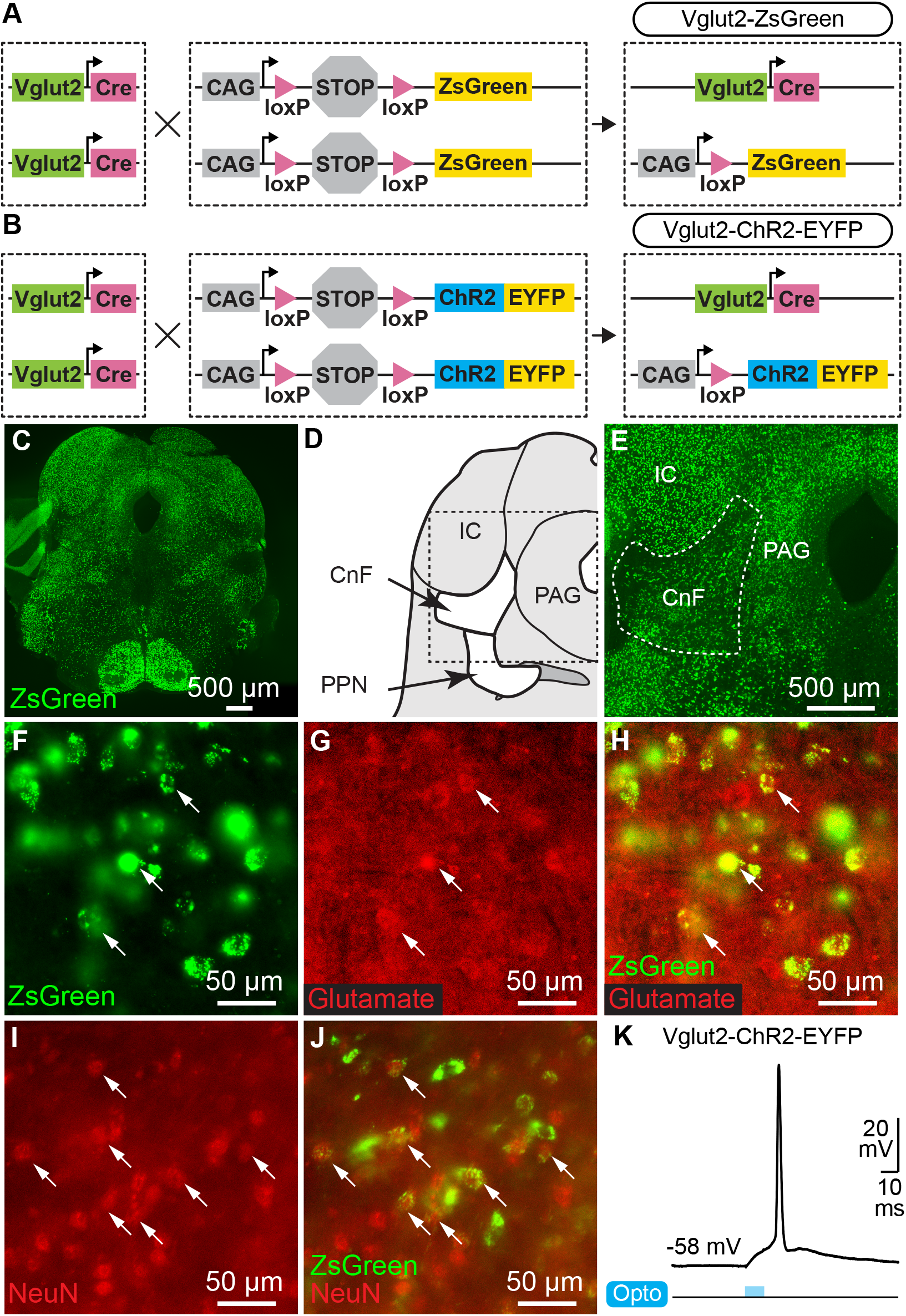
Cre-dependent expression of channelrhodopsin or ZsGreen in neurons of the Mesencephalic Locomotor Region (MLR) expressing the vesicular glutamate transporter 2 (Vglut2). **A.** For anatomical experiments, homozygous mice expressing Cre-recombinase under the control of the Vglut2 promoter (Vglut2-Cre, see Methods) were crossed with homozygous mice with ZsGreen preceded by a STOP cassette flanked by loxP sites preventing ZsGreen expression. In the resulting heterozygous mice (Vglut2-ZsGreen), if Vglut2 is expressed during cell lifetime, Cre-dependent recombination removes the STOP cassette, allowing permanent expression of ZsGreen under control of the CAG promoter. **B.** For optogenetic experiments, mice homozygous for Vglut2-Cre were crossed with mice homozygous for channelrhodopsin (ChR2) and enhanced yellow fluorescent protein (EYFP) preceded by a STOP cassette flanked by loxP sites preventing their expression. In the resulting heterozygous mice (Vglut2-ChR2-EYFP), if Vglut2 is expressed during cell lifetime, Cre-dependent recombination removes the STOP cassette, allowing permanent expression of ChR2-EYFP under control of the CAG promoter. **C.** Photomicrographs of transversal brain slices from Vglut2-ZsGreen mice at the MLR level. **D.** Schematic representation of a brain slice at the MLR level. **E.** Higher magnification of the brain slice in C at the level of the CnF. **F-H.** Epifluorescence images taken in the same field of view in the CnF, showing typical examples of cells expressing ZsGreen (green, F), cells immuno-positive for glutamate (red, G), and the two markers merged (H). **I-J.** Images taken in the CnF, showing that cells expressing ZsGreen (green) are immunopositive for the neuronal marker NeuN (red). **K.** Whole cell patch-clamp recording of a neuron recorded in a brainstem slice of a Vglut2-ChR2-EYFP mouse at the level of the MLR. The neuron spikes an action potential at short latency in response to a 10 ms blue light pulse. IC, inferior colliculus PAG, periaqueductal grey, PPN, pedunculopontine nucleus.

### Optical fiber implantation

Mice were anesthetized using isoflurane (induction: 5%, 500 mL/min; maintenance: 1.5-2.5%, 100 mL/min) delivered by a SomnoSuite (Kent Scientific, Torrington, CT, USA). Mice were placed in a Robot Stereotaxic instrument coupled with StereoDrive software (Neurostar, Tübingen, Germany) to perform unilateral implantation of an optical fiber (200 µm core, 0.22 NA, Thorlabs, Newton, NJ, USA) held in a 5 mm ceramic or stainless-steel ferrule 500 µm above the right CnF at −4.80 mm anteroposterior, +1.10 mm mediolateral, −2.40 mm dorsoventral relative to bregma [11, 12]. The ferrule was secured on the cranium using two 00-96 x 1/16 mounting screws (HRS scientific, QC, Canada) and dental cement (A-M Systems, Sequim, WA, USA).

### *In vivo* optogenetic stimulation

The optical fiber was connected using a pigtail rotary joint (Thorlabs) to a 470 nm laser (Ikecool, Anaheim, CA, USA) or a 589 nm laser (Laserglow, ON, Canada) driven by a Grass S88X that generated the stimulation trains (10 s train, 10 ms pulses, 20 Hz) [11, 12]. To visualize optogenetic stimulation, a small (diameter 0.5 cm), low-power (0.13 W) red LED that received a copy of the stimulation trains was placed in the camera’s field of view. The 470 nm light source was adjusted to 6-27% of laser power and the 589 nm to 40-53% of laser power. The corresponding power measured at fiber tip with a power meter (PM100USB, Thorlabs) was 0.1-16.0 mW for the 470 nm laser and 1.7-9.4 mW for the 589 nm laser.

### Open-field locomotion

Locomotor activity was filmed from above in a 40 × 40 cm arena at 30 fps using a Logitech Brio camera coupled to a computer equipped with ANY-maze software (Stoelting Co., Wood Dale, IL, USA) or using a Canon Vixia HF R800 camera. Locomotor activity was recorded during trials of 15 min during which 10 stimulation trains were delivered every 80 s at various laser powers. Video recordings were analyzed on a computer equipped with DeepLabCut (version 2.1.5.2), a software based on deep learning to track user-defined body parts [47, 48], and a custom Matlab script (Mathworks, Natick, MA, USA). We tracked frame by frame the body center position, the corners of the arena for distance calibration and the low-power LED to detect optogenetic stimulations. Timestamps were extracted using Video Frame Time Stamps (Matlab File Exchange). Body center positions and timestamps were used to calculate locomotor speed in cm/s. To compare and average speed over time for different stimulations, the data was downsampled to 20 Hz. Body center positions were excluded if their likelihood of detection by DeepLabCut was < 0.8, if they were outside of the open field area, or if body center speed exceeded the maximum locomotor speed recorded in mice (334 cm/s, [49]).

For offline analysis of turning movements in the arena’s corners, we defined regions of interest (ROIs) as circles (radius 20 cm) centered on each corner. The turning point was defined as the intersection of the mouse’s trajectory with the bisector of each corner (i.e. diagonal of the corner, Fig. 5D). The coordinates of the turning point were calculated using Curve intersections (Matlab File Exchange). Within an ROI, a turn was defined as a trajectory that started at least 5 cm away from the turning point, crossed the diagonal, and ended at least 5 cm away from the turning point. Turns were excluded if the mouse crossed the diagonal more than once without leaving the ROI. The turn angle was measured between the first point of the trajectory, the turning point, and the last point of the trajectory.

Locomotor speed during the start of the turn (“entry speed”), around the turning point (“turn speed”), and during the end of the turn (“exit speed”) were measured using the distance of each point of the trajectory to the turning point. These distance values were binned (width: 1 cm) and speed values were averaged per bin. Entry speed was averaged from the four most distal distance bins before the turning point and at least 5 cm away from the turning zone (2 cm radius around turning point). Turn speed was averaged from the four distance bins located within the turning zone. Exit speed was averaged from the four most distal distance bins after the turning point and at least 5 cm away from the turning zone. Turns were removed from the analysis if fewer than four bins were available to calculate entry, turn, or exit speed.

### Footfall patterns and limb kinematics

To label hindlimb joints for offline tracking with DeepLabCut, mice were anesthetized with isoflurane (induction: 5%, 500 mL/min; maintenance: 1.5-2.5%, 100 mL/min), the hindlimb was shaved and white dots (diameter ∼2 mm) were drawn on the iliac crest, hip, knee, ankle, and metatarsophalangeal (**MTP**) joints, and toe tip using a fine-tip, oil-based paint marker (Sharpie). For footfall pattern tracking, no labeling of paw underside was needed. Animals recovered for 20 min after anesthesia and were placed in a 1 m long, 8 cm wide transparent corridor. The footfall pattern and hindlimb kinematics were recorded at 300 fps using two high speed Genie Nano Camera M800 cameras (Teledyne DALSA, Waterloo, ON, Canada) coupled to a computer equipped with Norpix Streampix software (1st Vision, Andover, MA, USA). Hindlimb kinematics were recorded with a camera placed on the side of the corridor. Footfall patterns were recorded with a camera placed on the side and directed toward a 45-degree mirror placed below the corridor. For distance calibration, 4 markers (diameter 0.5 cm) were distributed 5 cm apart and placed in the field of view of each camera. To detect optogenetic stimulation, a low-power LED that received a copy of the stimulation trains was placed in the field of view of both cameras. Animals were recorded during spontaneous locomotion evoked by a gentle touch of the animal’s tail, and during optogenetic-evoked locomotion.

For the footfall pattern, videos recorded from below were used to track the position of the MTPs of the four paws with DeepLabCut [47, 48, 50]. Paw speeds were calculated and smoothened with a moving average (on 5 frames) using a custom Matlab script. Touchdown and lift-off were defined for each paw as the time points at which each MTP speed respectively fell below or rose above 15 cm/s. The touchdown and lift-off time points of each limb were identified using Curve intersections (Matlab File Exchange) and were normalized to the step cycle of the left hindlimb to generate normalized gait diagrams [11]. A step cycle was defined as the time between two consecutive touchdowns of the left hindlimb [12]. Cycle duration, stance phase duration, swing phase duration and stride length were calculated [12].

For hindlimb kinematics, the positions of the joints and toe tip were detected using DeepLabCut. The moving average of the MTP speed was used to determine the stance and swing phases by detecting the touchdown and lift-off times with a speed threshold of 9 cm/s. The joint positions were used to extract the angles of the hip, knee and ankle joints (Fig. 4C). The angular variations as a function of time were normalized to step cycle duration using MTP touchdown times as a reference [51].

Frames were excluded from the analysis if the MTPs of any paw (for the footfall pattern) or any limb joints or the toe tip (for limb kinematics) had a likelihood of detection < 0.8 by DeepLabCut. Frames were excluded from the analysis if any paw’s or joint’s speed exceeded 400 cm/s, i.e. the maximum locomotor speed of a mouse with a 20% margin to account for increased speed of individual body parts [49].

### Statistical analysis

Data are presented as mean ± standard error of the mean (**SEM**) unless stated otherwise. Statistical analyses were done using Sigma Plot 12.0. Normality was assessed using the Shapiro-Wilk test. Equal variance was assessed using the Levene test. Parametric analyses were used when assumptions for normality and equal variance were respected, otherwise non-parametric analyses were used. To compare the means between two dependent groups, a two-tailed paired-t test was used. For more than two dependent groups, a parametric one-way analysis of variance (ANOVA) for repeated measures or a non-parametric Friedman ANOVA for repeated measures on ranks was used. ANOVAs were followed by a Student Newman-Keuls post-hoc test for multiple comparisons between groups. Statistical differences were assumed to be significant when *P* < 0.05.

For genotyping, patch-clamp recordings, histology and immunofluorescence, patch-clamp recordings, specificity of the antibodies and of the transgenic mice, and details on DeepLabCut networks, please see the supplementary material.

## Results

We targeted glutamatergic cells that expressed Vglut2 in the CnF for optogenetic stimulation. We examined the presence of such cells by crossing mice expressing the Cre-recombinase under control of the *Vglut2* promoter (Vglut2-Cre mouse) with mice expressing a green fluorescent protein in a Cre-dependent manner (ZsGreen-lox mice) (Fig. 1A). In the offspring (Vglut2-ZsGreen mice), many cells were positive for ZsGreen in the CnF (n = 3 mice, Fig. 1C-E). Many of these cells were immunopositive for NeuN (n = 3 mice, Fig. 1I-J) and for glutamate (n = 3 mice, Fig. 1F-H). To stimulate these cells with blue light, we crossed Vglut2-Cre mice with mice expressing ChR2 in a Cre-dependent manner (ChR2-EYFP-lox) (Fig. 1B). Using patch-clamp recording in slices of the offspring (Vglut2-ChR2-EFYP mice), we validated that blue light elicited spiking at short latency in MLR neurons (n = 2 neurons from 2 mice, Fig 1K).

We then activated CnF neurons in freely moving Vglut2-ChR2-EYFP mice in an open-field arena (Fig. 2A). We implanted an optic fiber 500 µm above the right CnF and verified the implantation sites (n = 5 mice, Fig. 2B-C). Optogenetic stimulation of the CnF with blue light increased locomotor speed as shown by single animal data (Fig. 2D,F) and data pooled from 5 mice (Fig. 2H). Statistical analysis confirmed that speed was increased during optogenetic stimulation (*P* < 0.01 vs prestim, Student Newman-Keuls after a one-way ANOVA for repeated measures, *P* < 0.01) and decreased after light was switched off (*P* < 0.01 vs opto stim, Fig. 2J). Replacing the 470 with a 589 nm laser did not increase locomotion as shown by single animal data (Fig. 2E,G) and data pooled from 4 mice (*P* > 0.05 one-way ANOVA for repeated measures, Fig. 2I,K).

**Figure 2.**
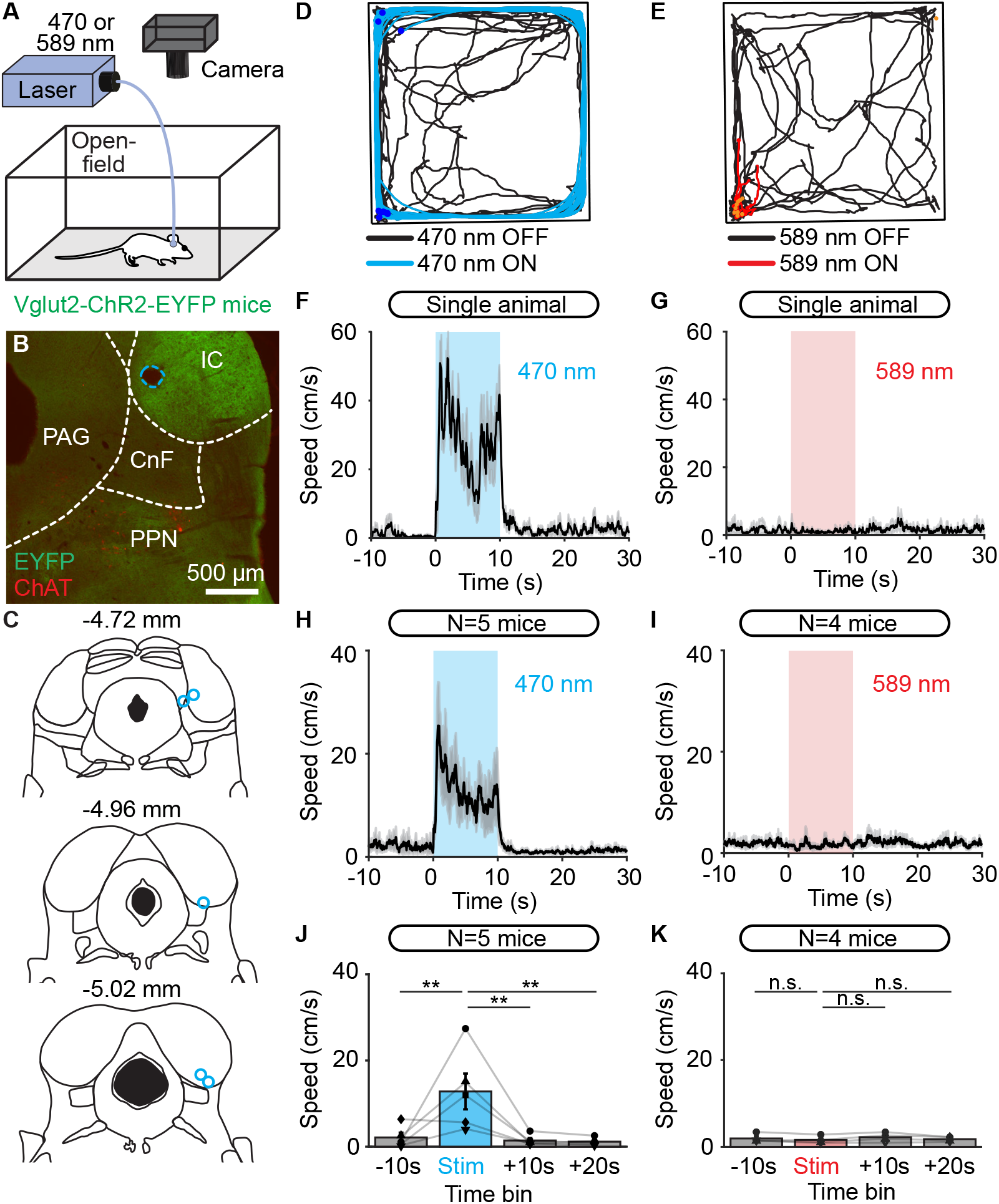
Optogenetic stimulation of the cuneiform nucleus (CnF) increases locomotor speed in the open field arena in Vglut2-ChR2-EYFP mice. **A.** An optic fiber implanted in the right CnF was connected to a blue laser (470 nm) or red laser (589 nm). Animals were placed in an open-field arena (40 × 40 cm) and their movements were recorded using a camera placed above. **B.** Photomicrograph showing the position of the optic fiber (dashed blue line) ∼500 μm above the target site. The cholinergic neurons of the pedunculopontine nucleus (PPN) (choline acetyltransferase positive, ChAT, red) and the expression of EYFP (green) are visible. **C.** Location of the optic fibers (blue circles) after histological verification as illustrated in B, with the relative position to the bregma. **D-E.** Raw data showing the effects of 10 optogenetic stimulations with a 470 nm laser (E, light blue lines, 10 s train, 20 Hz, 10 ms pulses, 11% of laser power) or a 589 nm laser (F, red lines, 10 s train, 20 Hz, 10 ms pulses, 53% of laser power). A time interval of 80 s was left between two trains of stimulation. The position of the animal’s body center was tracked frame by frame with DeepLabCut (see methods). Dark blue dots (D) and orange dots (E) illustrate the onset of each stimulation. **F-G.** Locomotor speed (mean ± sem) as a function of time before, during and after a 10 s optogenetic stimulation (onset at t = 0 s) with a 470 nm laser (G) or 589 nm laser in a single animal (same animal as in D,E). **H-I.** Locomotor speed (mean ± sem) before during and after optogenetic stimulation with a 470 nm laser in 5 animals (H, 10 stimulations per animal, 10-24% of laser power) and with the 589 nm laser in 4 animals (I, 10 stimulations per animal, 40-53% of laser power). **J-K.** Locomotor speed (mean ± sem) before (−10 to 0 s), during (0 to +10s), and after optogenetic stimulation (+10 to +20 s and +20 to +30 s) with the 470 nm laser in 5 animals (J) and with the 589 nm laser in 4 animals (K) (10 stimulations per animal, ** *P* < 0.01, n.s. not significant, *P* > 0.05, Student-Newman-Keuls test after a one way ANOVA for repeated measures, *P* < 0.01 in J and *P* > 0.05 in K).

Next, we compared spontaneous and optogenetic-evoked locomotion in a transparent linear corridor. We tracked the movements of each paw frame by frame using DeepLabCut [47, 48] (Fig. 3A). The footfall pattern was similar during spontaneous and optogenetic-evoked locomotion (Fig. 3B-C). We normalized the cycle duration as a function of the left hindlimb movements and observed again similar gait diagrams during spontaneous and optogenetic-evoked locomotion as shown by single animal data (Fig. 3D-E) and data pooled from 4 mice (Fig. 3F-G). We noticed, however, that mice were stepping faster during optogenetic-evoked locomotion as the cycle duration was shorter (*P* < 0.05 vs. spontaneous, paired test, n = 4 animals, Fig. 3H) while stride length did not differ (*P* > 0.05, Fig. 3I). This was associated with a shorter stance duration (*P* < 0.01, Fig. 3J), but no modification of swing duration (*P* > 0.05, Fig. 3K), consistent with the specific modulation of stance duration when speed increases during natural locomotion [52]. Altogether, this indicated that optogenetic CnF stimulation evoked a normal footfall pattern in Vglut2-ChR2-EYFP mice.

**Figure 3.**
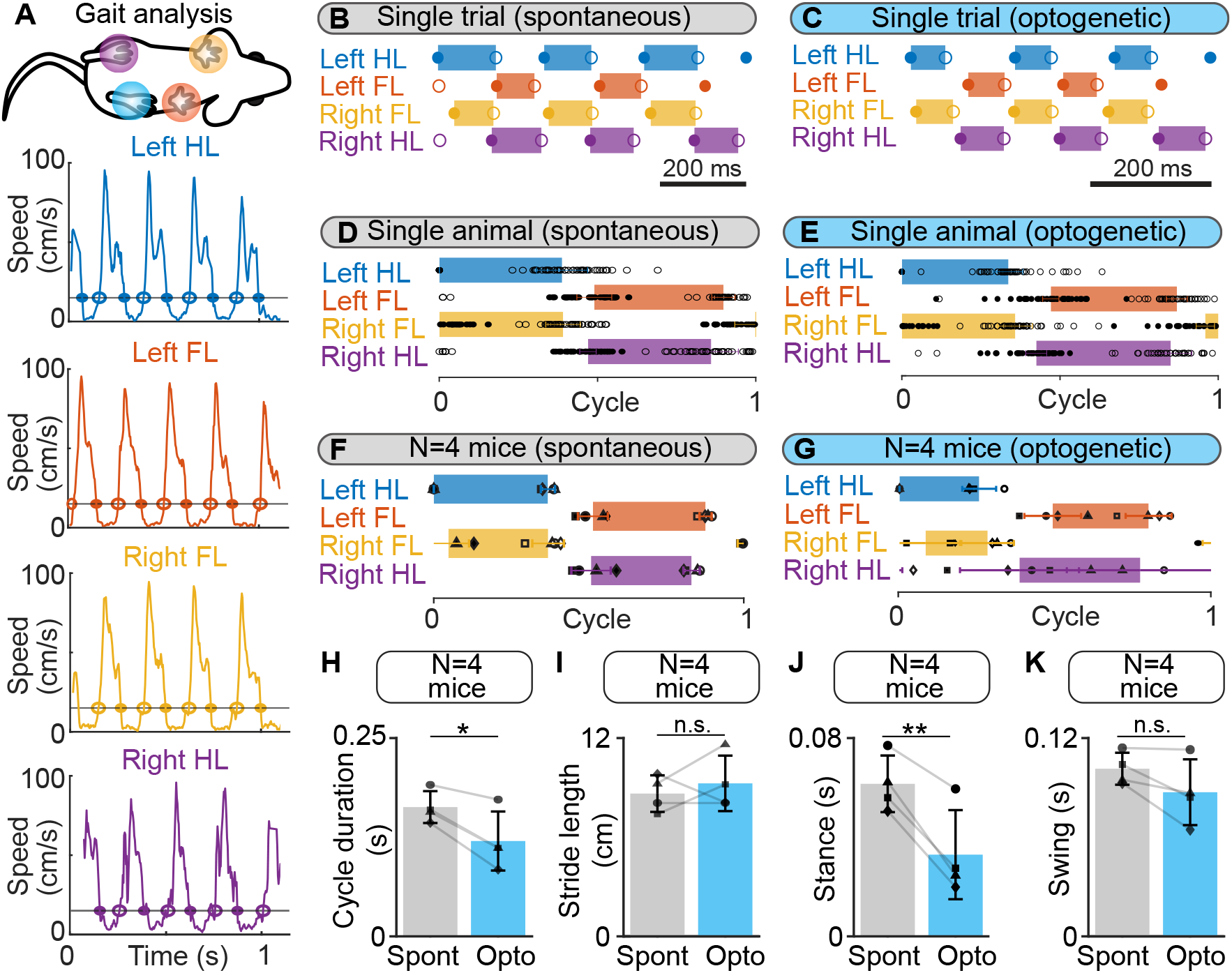
Gait diagrams during spontaneous locomotion and locomotion evoked by optogenetic stimulation of the cuneiform nucleus (CnF) in Vglut2-ChR2-EYFP mice in a linear corridor. **A.** Mouse forelimbs (FL) and hindlimbs (HL) were filmed from below at 300 fps in a transparent linear corridor and the position of each limb was tracked frame by frame with DeepLabCut (see methods). The four panels show the movement speed of each paw as a function of time. Cycle duration was defined as the time duration between two touchdowns of the left hindlimb (HL) using a speed threshold of 15 cm/s to define the transitions between swing and stance phases. Full circles are touchdowns; empty circles are lift-offs. **B-C.** Gait diagram for each limb obtained during a single spontaneous locomotor bout (B) and a locomotor bout evoked by optogenetic stimulation in the same animal (470 nm laser, 10 s train, 20 Hz, 10 ms pulses, 8 % of laser power). **D-E.** Gait diagrams during a normalized locomotor cycle, showing the stance phase start (mean ± SD) and end (mean ± SD) during 16 spontaneous locomotor bouts (45 steps) and during 8 locomotor bouts evoked by optogenetic stimulation in the same animal (48 steps, same stimulation parameters as in E). Cycle has been normalized to the left HL’s touchdown. **F-G.** Normalized gait diagram showing the touchdown (mean ± SD) and lift-off (mean ± SD) pooled from 4 mice during a total of 55 spontaneous locomotor bouts (8-16 trials per animal, 13-45 steps per animal) and from 4 mice during a total of 30 locomotor bouts (6-8 bouts per animal, 8-48 steps per animal) evoked by CnF optogenetic stimulation (470 nm laser, 10 s train, 20 Hz, 10 ms pulses, 8-15% of laser power). The data from each animal are illustrated with a different symbol. **H-K.** Comparison of cycle duration (H), stride length (I), stance duration (J) and swing duration (K) in 4 animals during spontaneous optogenetic-evoked locomotion (same animals as in F,G). \**P* < 0.05, \*\**P* < 0.01, n.s. not significant, *P* > 0.05, paired t tests).

We then compared the limb kinematics by tracking each hindlimb joint (iliac crest, hip, knee, ankle, MTP) and the toe tip using DeepLabCut (Fig. 4A). The stick diagrams were similar during spontaneous and optogenetic-evoked locomotion (Fig. 4B). We compared the angular variations of the hip, knee, ankle and MTP joints as a function of time (Fig. 4C) and cycle duration was normalized relative to MTP movements (Fig. 4D). The angular variations were similar during spontaneous and optogenetic-evoked locomotion as shown by single animal data and data pooled from 4 mice (Fig. 4E). Statistical analysis revealed no difference of the angle amplitude of the four joints between the two conditions (*P* > 0.05 vs spontaneous, paired tests, n = 4 animals, Fig. 4F). Altogether, this indicated that optogenetic CnF stimulation evoked normal limb kinematics.

**Figure 4.**
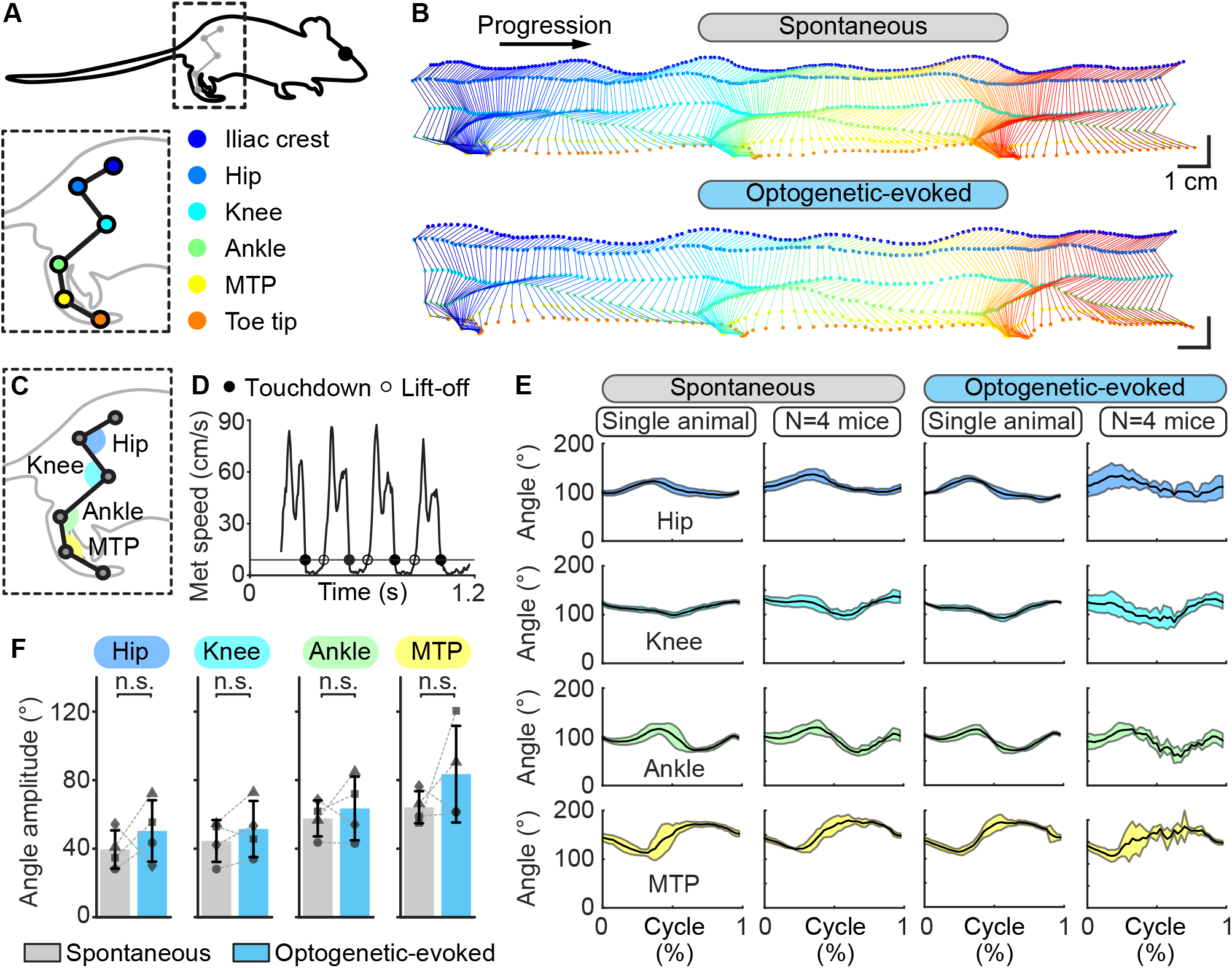
Hindlimb kinematics during spontaneous locomotion and locomotion evoked by optogenetic stimulation of the cuneiform nucleus (CnF) in Vglut2-ChR2-EYFP mice in a linear corridor. **A.** Six hindlimb joints were labeled with a white paint marker and were filmed from the side at 300 fps in a transparent linear corridor and the trajectory of each joint was extracted with DeepLabCut and plotted in a different color (see methods). **B.** Side view of the hindlimb joints during spontaneous locomotion (top) and optogenetic-evoked locomotion (470 nm laser, 10 s train, 20 Hz, 10 ms pulses, 8% of laser power) (bottom). Total time elapsed from first to last frames is 700 ms (top) and 500 ms (bottom). **C.** Joint angles at the hip, knee, ankle and metatarsophalangeal joint (MTP) levels were calculated frame by frame using the position of the joint of interest and those of two proximal joints. **D.** Cycle duration was defined as the time duration between two consecutive touchdowns of the MTP using a speed threshold of 9 cm/s to define the transitions between swing and stance phases. Full circles are touchdowns; empty circles are lift-offs. **E.** Joint angles (mean ± SD) at the hip, knee, ankle and MTP levels plotted for a normalized locomotor cycle during spontaneous locomotion (29 steps) and locomotion evoked by optogenetic stimulation (31 steps) (470 nm laser, 10 s train, 20 Hz, 10 ms pulses, 8% of laser power). For the pooled data, joint angles (mean ± SD) of 4 animals plotted for a normalized locomotor cycle during spontaneous locomotion (12-29 steps per animal) and locomotion evoked by optogenetic stimulation are shown (1-31 steps per animal) (470 nm laser, 10 s train, 20 Hz, 10 ms pulses, 8-15% of laser power). **F.** Amplitude of the hip, knee, ankle and MTP angles measured in 4 animals during spontaneous and optogenetic evoked locomotion (same data as in E). n.s., not significant, *P* > 0.05, paired t tests.

We then examined whether freely behaving mice could brake or turn during optogenetic CnF stimulation in the open-field arena. Inspection of the speed as a function of time uncovered oscillations during optogenetic stimulation (Fig. 5A-B). We plotted the speed as a function of the location of the animal in the arena and found that speed decreased in the corners of the arena, where the animal was performing turning movements (Fig. 5C). This suggested that during CnF stimulation, the animal dynamically controlled speed as a function of environmental cues. We furthered studied this phenomenon by analyzing locomotor movements in each corner of the arena during CnF stimulation (Fig. 5D). We defined ROIs as circles (20 cm radius) centered on each corner. The trajectories of a single mouse within the four ROIs during CnF stimulation are illustrated in Fig. 5E. We defined the turning point as the intersection between the mouse’s trajectory and the corner’s bisector (i.e. arena’s diagonal). Plotting the speed relative to the distance from the turning point indicated that speed was lower around the turning point in single animal data (Fig. 5G) as in data pooled from 5 mice (Fig. 5F,H). Statistical analysis showed that speed decreased by ∼61% during the turn (*P* < 0.05 vs. entry speed, Student Newman-Keuls after a one-way ANOVA for repeated measures, *P* < 0.01, Fig. 5I). Speed increased when exiting the corner (*P* < 0.01 vs. turn speed, Fig. 5I) to values that were not different from the entry speed (*P* > 0.05 vs entry speed, Fig. 5I). This indicated that the slowdown was transient and linked to the turn, after which ongoing MLR stimulation regained control over speed.

**Figure 5.**
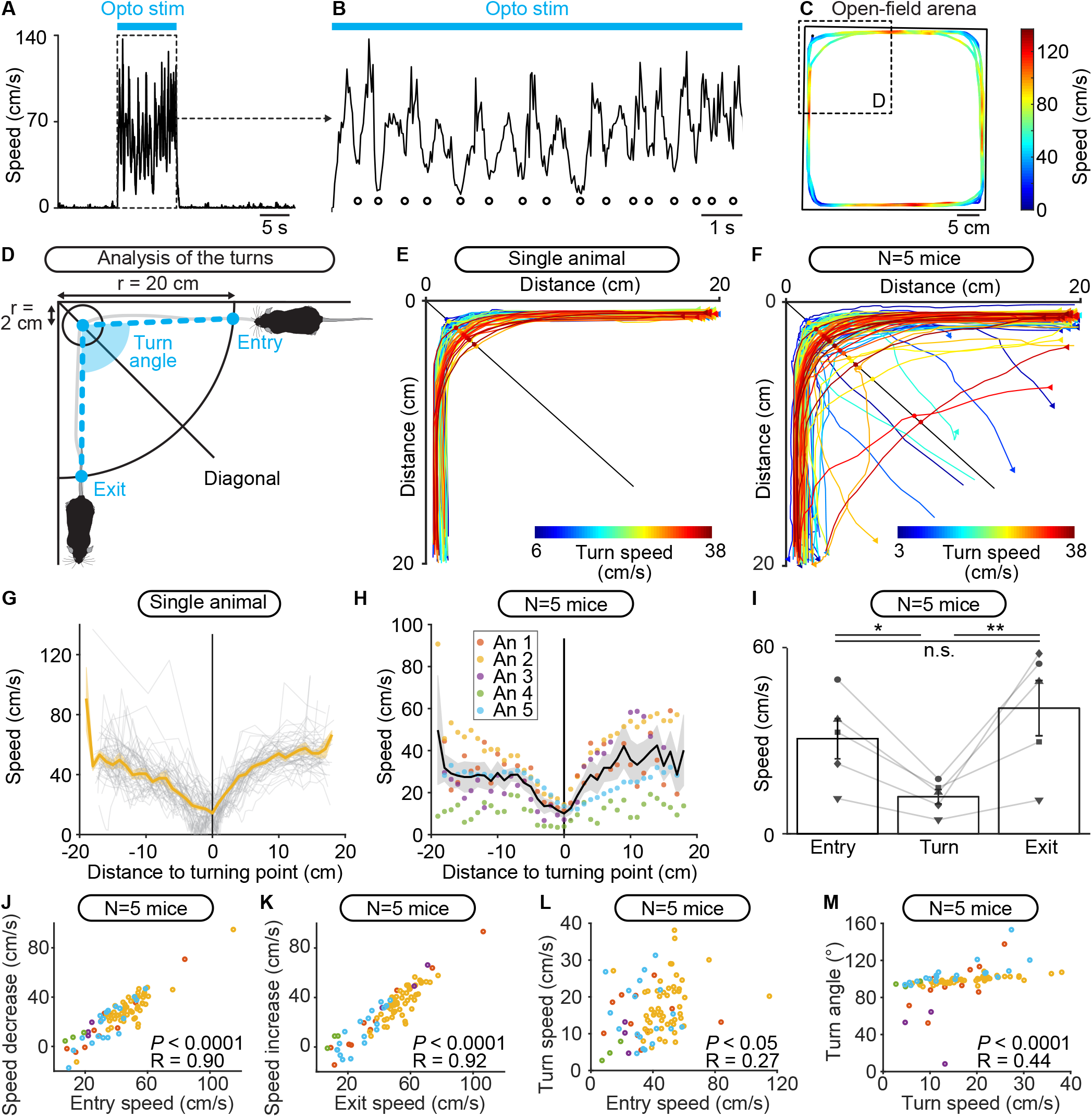
Freely behaving mice brake and turn during locomotion evoked by optogenetic stimulation of the cuneiform nucleus (CnF) in Vglut2-CHR2-EYFP mice in the open field arena. **A.** Raw data showing that locomotor speed is modulated during optogenetic stimulation of the CnF (470 nm laser, 10 s train, 20 Hz, 10 ms pulses, 11% of laser power). **B**. Magnification of A showing rhythmic speed decrease (white circles) during CnF stimulation (blue solid line). **C.** Color plot illustrating the locations of the speed decreases during CnF stimulation in the open-field arena (same data as in A-B). Colder colors (blue) illustrate slower speeds. **D.** Animal’s speed was measured when moving in 20 cm circles centered on each corner of the arena. The speed at the entry of the corner, during the turn (in a 2 cm circle centered on the location where the animal crossed the corner’s diagonal) and at the exit of the corner were calculated (see methods). The turn angle was measured between the positions of the animal i) at the entry of the corner, ii) during the turn, and iii) when exiting the corner (see methods). **E-F.** Raw data showing the extracted locomotor trajectories in the corners of the arena during optogenetic-evoked locomotion in a single animal (E, 470 nm laser, 10 s train, 20 Hz, 10 ms pulses, 11% of laser power) and in 5 animals (F, 470 nm laser, 10 s train, 20 Hz, 10 ms pulses, 10-24% of laser power). Warmer colors (red) illustrate trajectories with higher speeds during the turn. Triangles illustrate movement onsets. Dots illustrate diagonal crossings. **G.** Locomotor speed as a function of the distance to the corner’s diagonal during each turn (grey) shown in E for a single animal. In orange, the averaged speed (± sem) is shown. **H.** Locomotor speed as a function of the distance to the diagonal during the turns of the 5 animals (An) illustrated in F. In black, the mean speed (± sem in grey) is shown. **I.** Entry speed, turn speed and exit speed for 5 animals (10 stimulations per animal, \**P* < 0.05, \*\**P* < 0.01, n.s., not significant, *P* > 0.05, Student-Newman-Keuls test after a one way ANOVA for repeated measures, *P* < 0.01). **J.** Relationship between the speed at corner entry, and the difference between entry speed and turn speed in 5 mice (linear fit, *P* < 0.0001, R = 0.90, n = 87 turns pooled from 50 stimulations, 10 stimulations per animal). **K.** Relationship between the speed at corner exit, and the difference between exit speed and turn speed (linear fit, *P* < 0.0001, R = 0.92). **L.** Relationship between the speed at corner entry and the turn speed (linear fit, *P* < 0.05, R = 0.27). **M.** Relationship between the turn speed and the turn angle (linear fit, *P* < 0.0001, R = 0.44).

We examined the relationships between speed, braking and turn angle. We found a strong positive linear relationship between the entry speed, and the speed decrease between entry and turning zone in 5 mice (*P* < 0.0001, R = 0.90, Fig. 5J), and a strong positive linear relationship between the exit speed, and the speed increase between turning zone and exit (*P* < 0.0001, R = 0.92, Fig. 5K). We found a weak but significant positive linear relationship between entry speed and turn speed (*P* < 0.05, R = 0.27, Fig. 5L) and a positive linear relationship between turn speed and turn angle (*P* < 0.0001, R = 0.44, Fig. 5M). This last relationship is visible when looking at the trajectories color coded as a function of turn speed (Fig. 5E-F). This indicated that during CnF stimulation, wider angles were easier to negotiate at high speed than sharp ones, as reported during natural locomotion in mammals [1].

We examined the robustness of such scaled control of speed during turning when increasing CnF stimulation. Increasing the laser power applied to the CnF increased locomotor speed as shown by single animal data (Fig. 6A-B) and data pooled from 5 mice (Fig. 6C). We expressed the laser power and speed as a function of their maximal values per animal, and we found a strong positive sigmoidal relationship between laser power and speed (*P* < 0.01, R = 0.99, Fig. 6D). Such precise control of speed confirmed that we successfully targeted the CnF (Fig. 2C). Mice made to walk at increasing speeds imposed by increasing CnF stimulation were able to maintain successful braking and turning as shown by single animal data (Fig. 6E-F) and data pooled from 5 mice (Fig. 6G-H). The relationships describing the scaling of speed relative to the turn properties were conserved within this range of speeds (Fig. 6I-L). This indicated that CnF stimulation controls locomotor speed, without preventing the animal from precisely regulating braking and turning, likely through dynamic integration of environmental cues.

**Figure 6.**
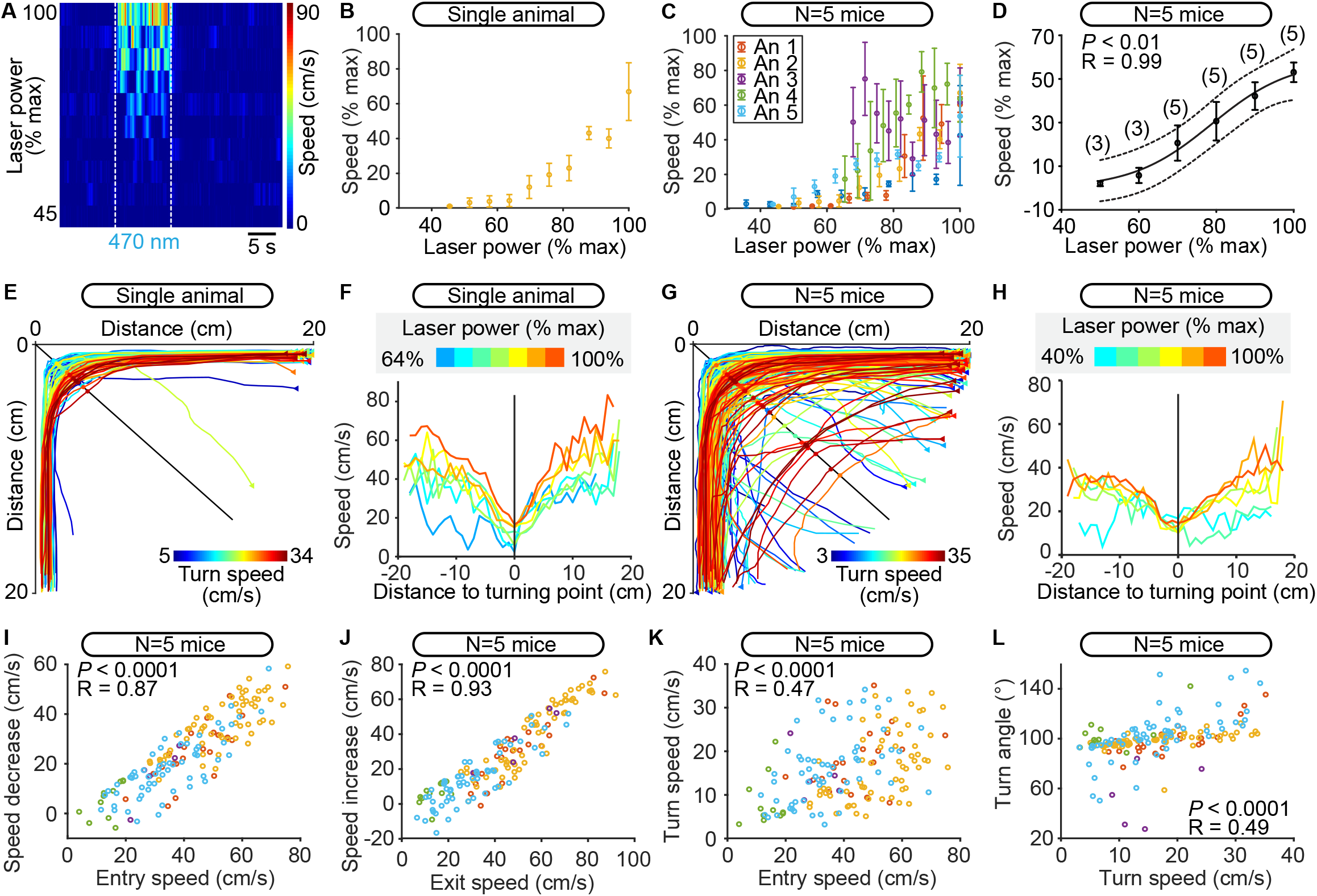
Robustness of scaled control of speed during turning at different speeds controlled by the level of optogenetic stimulation of the cuneiform nucleus (CnF) in Vglut2-ChR2-EYFP mice in the open field arena. **A.** Color plot illustrating the increase in locomotor speed (cm/s) evoked by increases in laser power (6-13%) in a single animal (470 nm laser, 10 s train, 20 Hz, 10 ms pulses). Each line illustrates the speed as a function of time for a given laser power expressed as a percentage of maximal laser power used for this animal. White dotted lines indicate the onset and offset of optogenetic stimulation. Warmer colors (red) indicate higher speeds. **B.** Locomotor speed (1-42 cm/s) as a function of laser power (6-13%) in one animal. Each dot represents the speed (mean ± sem) over 3 stimulations. Speed and laser power were expressed as a function of their maximal values. **C.** Relationship between locomotor speed (0.2-42.0 cm/s) and increasing laser power (6-27%) for all animals. Data from each mouse are illustrated with a different color. Each dot represents the speed (mean ± sem) over 3 stimulations. Speed and laser power were expressed as a percentage of their maximal values per animal. **D.** Relationship between locomotor speed (mean ± sem) and laser power in the same animals as in C, this time with data binned as a function of laser power used per animal with a bin size of 10%. The data followed a sigmoidal function (solid black line, *P* < 0.01, R = 0.99). The dotted lines illustrate the 95% prediction intervals. **E, G.** Raw data showing the extracted locomotor trajectories in the corners of the arena during optogenetic-evoked locomotion in a single animal (E) and in 5 animals (G). Triangles illustrate movement onsets. Dots illustrate diagonal crossings. Warmer colors (red) illustrate higher turn speeds. **F, H.** Locomotor speed as a function of the distance to the diagonal during the turns for increasing power of optogenetic stimulation of the CnF in a single animal (F, 470 nm laser, 10 s train, 20 Hz, 10 ms pulses, 6-13% of laser power) and in 5 animals (H, 470 nm laser, 10 s train, 20 Hz, 10 ms pulses, 6-27% of laser power). Warmer colors (red) indicate stronger optogenetic stimulation of the CnF. Laser powers were normalized as a percentage of their maximal value used per animal and were binned in H (bin width: 10%). In F, each curve was obtained from 1-14 turns in a single animal. In H, each curve was obtained from 2-73 turns pooled from 5 animals. **I.** Relationship between the speed at corner entry, and the difference between entry speed and turn speed (linear fit, *P* < 0.0001, R = 0.87, N = 157 turns pooled from 150 stimulations, 30 stimulations per animal). **J.** Relationship between the speed at corner exit, and the difference between exit speed and turn speed (linear fit, *P* < 0.0001, R = 0.93). **K.** Relationship between the speed at corner entry and the turn speed (linear fit, *P* < 0.0001, R = 0.47). **L.** Relationship between the turn speed and the turn angle (linear fit, *P* < 0.0001, R = 0.49).

## Discussion

We show in Vglut2-ChR2-EYFP mice that optogenetic stimulation of the CnF with blue light evoked locomotion and that increasing laser power increased speed. Replacing the blue laser with a red laser evoked no locomotion. In a linear corridor, footfall patterns and limb kinematics were largely similar during spontaneous and optogenetic-evoked locomotion. In the open-field arena, mice could brake and perform sharp turns (∼90⁰) when approaching a corner during CnF stimulation. Speed decrease during the turn was scaled to speed before the turn, and turn speed was scaled to turn angle. We verified the stimulation sites in the CnF and showed that many Vglut2-ZsGreen cells in the CnF were positive for NeuN and glutamate. Using patch-clamp recordings in brainstem slices we showed that blue light evoked short latency spiking. Altogether, our study indicates that the CnF controls locomotor speed without preventing the animal from integrating environmental cues to perform braking and turning movements, and thereby smoothly navigate the environment.

### Methodological considerations

We cannot exclude that some non-glutamatergic neurons were stimulated in the CnF of Vglut2-ChR2-EYFP mice. We crossed Vglut2-Cre with ChR2-EYFP-lox mice. In the offspring (Vglut2-ChR2-EYFP), if Vglut2 is expressed during cell lifetime, ChR2 is expressed permanently under control of the CAG promoter even if Vglut2 is not expressed anymore [53, 54]. Mainly glutamatergic and GABAergic neurons are present in the CnF (for review [3]). Although the expression of Vglut2 was detected in some GABAergic neurons in the mammalian brain (for review [55]), it is unlikely that we stimulated GABAergic neurons. Two arguments indicating that we successfully targeted CnF glutamatergic neurons are the normal gait diagrams and limb kinematics, and the precise control of speed when increasing laser power. These effects are consistent with results obtained in Vglut2-Cre mice optogenetically stimulated in the CnF following injection of an AAV encoding for ChR2 in a Cre-dependent manner [11, 12].

### Brainstem control of speed

Our results support the idea that MLR glutamatergic neurons play a key role in the initiation of forward symmetrical locomotion and in the control of speed by sending input to reticulospinal neurons (lamprey [4, 3], salamander [7, 18], mouse [20, 8, 9, 10, 11, 12]) that send input to excitatory neurons of the locomotor Central Pattern Generator (lamprey [14], zebrafish [16, 17], mouse: [19, 10], for review [56]). In mice, the reticulospinal neurons relaying MLR locomotor commands are in the LPGi [10].

In addition, we show that MLR stimulation does not prevent mice from braking and turning following integration of environmental cues. The turning and braking movements recorded here displayed the same characteristics as those shown by mammals during natural locomotion. At high speed, mice used fewer sharp turns (i.e. higher angles), consistent with observations in wild northern quolls, which reduce their locomotor speed more during turns with a smaller radius [1].

### Brainstem control of braking and turning

Our observations support the idea that distinct reticulospinal neurons control speed and turning/braking movements [29, 30, 31]. First, our data indicate that a substrate for turning movements is activated transiently during MLR stimulation when approaching a corner. This motor signature closely matches that previously recorded when selectively activating the brainstem circuit for turning [30, 31]. Indeed, unilateral activation of Chx10-positive reticulospinal neurons in the Gi produces an ipsilateral turn together with a decrease in speed [30, 31]. Gi-Chx10 reticulospinal neurons receive no input from the MLR, but a major input from the contralateral superior colliculus (SC), a region involved in visuomotor transformations [30]. Such connectivity is relevant to the behavioral task mice had to solve here in the open-field arena, i.e. integrating visual cues to avoid the arena’s corner during locomotion evoked by MLR stimulation (Fig. 7). Altogether, this suggests that the brainstem substrates for braking and turning ([29, 30, 31], see also [57]) can be recruited during MLR stimulation, therefore allowing the animal to smoothly navigate the environment (Fig. 7).

**Figure 7.**
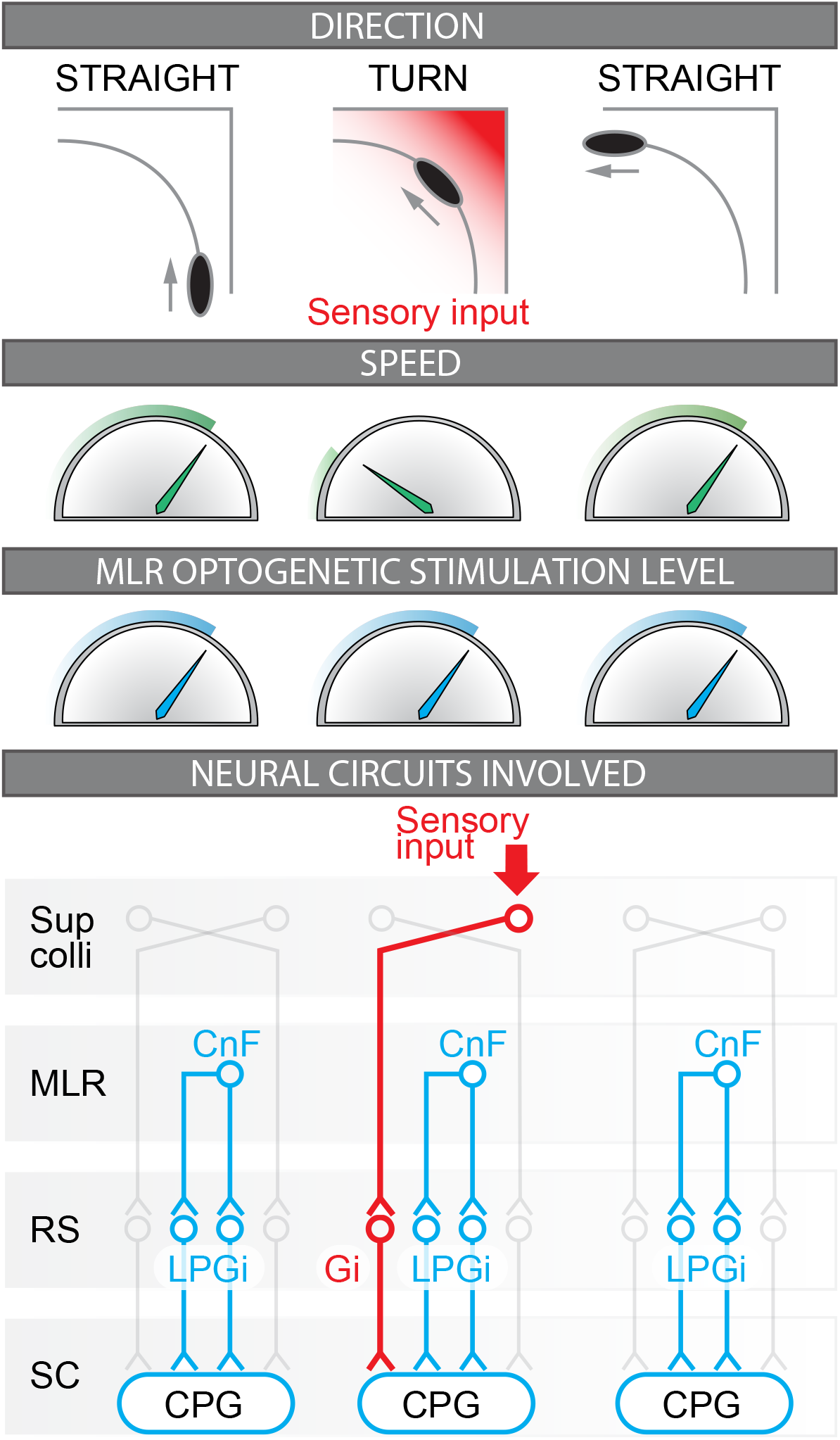
Coordinated control of locomotor speed and turning movements during optogenetic stimulation of the Mesencephalic Locomotor Region (MLR). Illustration of the relation between motion direction, locomotor speed, and MLR optogenetic stimulation level observed during the present study. The neural circuits likely involved before, during, and after the turn are illustrated. Before the turn, forward locomotion is evoked by unilateral stimulation of glutamatergic neurons of the cuneiform nucleus (CnF, part of the MLR) that provides bilateral activation of reticulospinal (RS) neurons located in the lateral paragigantocellular nucleus (LPGi) that project to the spinal neurons of the central pattern generator (CPG) for locomotion (mouse [20, 8, 9, 10, 11, 12]. Also see corresponding studies in lamprey [5, 6, 14, 15], zebrafish [16, 17], and salamander [7, 18]). During the turn, the visual inputs conveying the approach of the corner are relayed by the superior colliculus (Sup colli) that sends projections to contralateral reticulospinal neurons of the gigantocellularis nucleus (Gi) that evoke ipsilateral braking and turning movements [29, 30, 31]; also see studies on the role of reticulospinal neurons in steering control in lamprey [21, 22, 23, 24], zebrafish [25, 26], salamander [27], and rat [28]). After the turn, the sensory inputs generated by the corner disappear, Gi neurons are deactivated, speed increases back to the value set by the steady MLR command, and forward symmetrical locomotion is restored.

Interestingly, the speed decreased to zero during some turns, i.e. mice transiently halted during optogenetic MLR stimulation (Fig. 5G). Future studies should examine which level of the locomotor circuitry is involved in this effect. At the reticulospinal level, “stop cells” could increase their activity to stop locomotion as shown in in basal vertebrates (lamprey [58, 59]). In mammals, halt is induced by a bilateral recruitment of Gi Chx10-positive reticulospinal neurons, i.e. the same neurons that induce turning when activated unilaterally [29, 30, 31] (see also [57]). Locomotor pauses and rhythm resetting were also reported when photoactivating Vglut2-positive neurons in the Gi in mice [60]. At the MLR level, local GABAergic neurons could stop locomotion likely by inhibiting MLR glutamatergic neurons [9, 12]. In lamprey, stimulation of the MLR, at lower stimulation intensity values than the ones evoking locomotion, stop locomotion by recruiting reticulospinal stop cells [59]. Two incoming inputs to the MLR could be involved. A transient increase in the GABAergic tone from the output stations of the basal ganglia could stop locomotion (lamprey [61, 62], mouse [63, 9]). Alternatively, increased activity from the output station of the basolateral amygdala could be involved, since activation of this region is synchronized with locomotor arrests in familiar places during exploratory behavior ([64], for review [65]).

### Conclusions

We show that optogenetic stimulation of the CnF in Vglut2-ChR2-EYFP mice controls locomotor speed without preventing braking and turning movements following integration of environmental cues. This supports the idea that distinct brainstem circuits control speed (“MLR-LPGi pathway”, [8, 9, 10, 11, 12]) and braking/turning movements in mammals (“SC-Gi pathway”, [29, 30, 31]) (Fig. 7). This also suggests that CnF glutamatergic neurons are a relevant target to improve navigation adaptable to the environment in conditions where locomotion is impaired such as Parkinson’s disease [37, 38, 39, 40] spinal cord injury [41, 42] and stroke [43] (for review [44]).

## ACKNOWLEDGMENTS

We thank Jean Lainé for technical assistance with the microscopy platform, Florian Bentzinger for providing access to the genotyping equipment.

## Author contributions

**CIvdZ:** Conceptualization; Data curation; Formal analysis; Investigation; Methodology; Software; Validation; Visualization; Roles/Writing - original draft; Writing - review & editing. **JB:** Data curation; Investigation; Methodology; Validation; Visualization; Writing - review & editing. **MF:** Data curation; Investigation; Methodology; Validation; Visualization; Writing - review & editing. **AF:** Data curation; Investigation; Methodology; Validation; Visualization; Writing - review & editing. **MV:** Methodology; Writing - review & editing. **AS:** Methodology; Software; Writing - review & editing. **TA:** Methodology; Software; Writing - review & editing. **PS:** Methodology; Resources; Writing - review & editing. **DR:** Conceptualization; Data curation; Formal analysis; Funding acquisition; Methodology; Project administration; Resources; Supervision; Validation; Visualization; Roles/Writing - original draft; Writing - review & editing.

## Supplementary material

### Genotyping

Mice were genotyped as previously described [1]. Briefly, DNA was extracted from ear punches using the Taq DNA polymerase (NEB, Ipswich, MA, United States). Genotyping was performed using the primers recommended by the supplier (Jackson laboratory). Vglut2-ires-Cre mice were genotyped using mixed primer PCR employing Vglut2-ires-Cre-Com-F (AAGAAGGTGCGCAAGACG), Vglut2-ires-Cre-Wt-R (CTGCCACAGATTGCACTTGA) and Vglut2-ires-Cre-Mut-R (ACACCGGCCTTATTCCAAG). Amplification of wild-type genomic DNA yielded a 245 bp PCR product whereas amplification from the mutant locus yielded a 124 bp PCR product. ChR2-lox mice and ZsGreen-lox mice were genotyped using mixed primer PCR employing ZsGreen-ChR2-lox-Wt-F (AAGGGAGCTGCAGTGGAG TA), ZsGreen-ChR2-lox-Wt-R (CCGAAAATCTGTGGGAAGTC), ZsGreen-ChR2-lox-Mut-R (GGCATTAAAGCAGCGTATCC), and either ChR2-lox-Mut-F (ACATGGTCCTGCTGGAGTTC) or ZsGreen-lox-Mut-F (AACCAGAAGTGGCACCTGAC). Amplification of wild-type genomic DNA yielded a 297 bp PCR product whereas amplification from the mutant ChR2-lox locus yielded a 212 bp PCR product and amplification of the mutant ZsGreen-lox locus yielded a 199 bp PCR product.

### Patch-clamp recordings

Coronal brainstem slices were obtained from 15-23-day old mice as previously described [2]. Briefly, mice were anesthetized with isoflurane (0.5-1 mL of isoflurane in a 1.5 L induction chamber) and decapitated with a guillotine. The cranium was opened and the brain removed to be dipped in an ice-cold sucrose-based solution (in mM: 3 KCl, 1.25 KH_2_PO4, 4 MgSO_4_, 26 NaHCO_3_, 10 Dextrose, 0.2 CaCl_2_, 219 Sucrose, pH 7.3–7.4, 300-320 mOsmol/kg) bubbled with 95% O_2_ and 5% CO_2_. MLR slices (350 μm thick) were prepared with a VT1000S vibrating blade microtome (Leica Microsystems, Concord, ON, Canada) and stored at room temperature for 1 hour in artificial cerebrospinal fluid (**aCSF**) (in mM: 124 NaCl, 3 KCl, 1.25 KH_2_PO_4_, 1.3 MgSO_4_, 26 NaHCO_3_, 10 Dextrose, and 1.2 CaCl_2_, pH 7.3–7.4, 290–300 mOsmol/kg) bubbled with 95% O_2_ and 5% CO_2_. Whole-cell patch-clamp recordings were done in a chamber perfused with bubbled aCSF under an Axio Examiner Z1 epifluorescent microscope (Zeiss, Toronto ON Canada), differential interference contrast (DIC) components, and an ORCA-Flash 4.0 Digital CMOS Camera V3 (Hamamatsu Photonics, Hamamatsu, Japan). Patch pipettes were pulled from borosilicate glass capillaries (1.0 mm outside diameter, 0.58 mm inside diameter; 1B100F-4, World Precision Instruments, FL, USA) using a P-1000 puller (Sutter Instruments). Pipettes with a resistance of 6–12 MΩ were filled with a solution containing (in mM) 140 K-gluconate, 5 NaCl, 2 MgCl_2_, 10 HEPES, 0.5 EGTA, 2 Tris ATP salt, 0.4 Tris GTP salt, pH 7.2–7.3, 280–300 mOsmol/kg, 0.05 Alexa Fluor 594 or 488, and 0.2% biocytin). Positive pressure was applied through the glass pipette and neurons were approached using a motorized micromanipulator (Sutter instruments). A gigaseal was established and the membrane potential was held at −60 mV. The membrane patch was suctioned, and the pipette resistance and capacitance were compensated electronically. Neurons were discarded when action potentials were less than 40 mV or when the resting membrane potential was too depolarized (>-45 mV). Patch-clamp signals were acquired with a Multiclamp 700B coupled to a Digidata 1550B and a computer equipped with PClamp 10 software (Molecular Devices, Sunnyvale, CA, USA). Optogenetic stimulations (475 nm, 10 ms pulses, 2.5-5% of LED power) were applied using the 475 nm LED of a Colibri 7 illumination system (Zeiss).

### Histology and immunofluorescence

Procedures were as previously reported [1]. Briefly, mice were anaesthetized using isoflurane (5%, 2.5 L per minute) and transcardially perfused with 50 mL of a phosphate buffer solution (0.1M) containing 0.9% NaCl (PBS, pH = 7.4), followed by 40 to 75 mL of a PBS solution containing 4% (wt/vol) of paraformaldehyde (PFA 4%). Post-fixation of the brains was performed in a solution of PFA 4% for 24 h at 4°C. Then, the brains were incubated in a PBS solution containing 20% (wt/vol) sucrose for 24 h before histology. Brains were snap frozen in methylbutane (−45°C ± 5°C) and sectioned at −20°C in 40 µm-thick coronal slices using a cryostat (Leica CM 1860 UV). Floating sections of the MLR were collected under a Stemi 305 stereomicroscope (Zeiss) and identified using the atlas of Franklin and Paxinos (2008) [3].

For immunofluorescence experiments, all steps were carried out at room temperature unless stated otherwise. The sections were rinsed in PBS for 10 min three times and incubated for 1h in a blocking solution containing 5% (vol/vol) of normal donkey serum and 0.3% Triton X-100 in PBS. The sections were then incubated at 4°C for 48 h in a PBS solution containing the primary antibody against choline acetyltransferase (ChAT) (goat anti-choline acetyltransferase, Sigma AB144P, lot 3018862 (1:100), RRID AB_2079751), the neuronal marker NeuN (rabbit anti-NeuN, Abcam AB177487, lot GR3250076-6 (1:1,000), RRID AB_2532109) or glutamate (rabbit anti-glutamate, Sigma G6642, lot 079M4802V (1:3,000), RRID AB_259946) and agitated with an orbital shaker. The sections were washed three times in PBS and incubated for 4 h in a solution containing the appropriate secondary antibody to reveal ChAT (donkey anti-goat Alexa 594, Invitrogen A11058, lot 1975275 (1:400), RRID AB_2534105) NeuN or glutamate (with a donkey anti-rabbit Alexa Fluor 594, Invitrogen A21207 lot 1890862 (1:400), RRID: AB_141637; or a donkey anti-rabbit Alexa Fluor 647, ThermoFisher A31573 lot 2083195 (1:400), RRID: AB_2536183). The slices were rinsed three times in PBS for 10 min and mounted on Colorfrost Plus glass slides (Fisher) with a medium with DAPI (Vectashield H-1200) or without DAPI (Vectashield H-1000), covered with a 1.5 type glass coverslip and stored at 4°C before observation. Brain sections were observed using a Zeiss AxioImager M2 microscope equipped with StereoInvestigator 2018 software (v1.1, MBF Bioscience). Composite images were assembled using StereoInvestigator. The levels were uniformly adjusted in Photoshop CS6 (Adobe) to make all fluorophores visible and avoid pixel saturation, and digital images were merged.

### Specificity of the antibodies

The AB177487 anti-NeuN has been widely used to label the neuronal marker NeuN (also called Fox-3, see [4, 5]) in mouse brain tissues by us [1] and others [6, 7]. According to the supplier, this monoclonal purified antibody (clone EPR12763) is directed towards a synthetic peptide of the residues 1-100 of the human NeuN. NeuN is present in most mouse neurons, but not in cerebellar Purkinje cells, olfactory bulb mitral cells, and retinal photoreceptor cells [4]. According to the supplier, AB177487 labels NeuN in HeLa cell lysates and in brains of mice, rats and humans. It detects two bands at 45-50 kDA in Western blots performed on mouse, rat or human brain tissues.

The G6642 anti-glutamate polyclonal antibody was used to label glutamatergic neurons in zebrafish, rat and mouse (see below). The dot-blot immunoassays carried out by the supplier and by Terada et al. (2009) [8] indicated that this antiserum recognizes L-glutamate, glutamate conjugated to keyhole limpet hemocyanin (KLH), glutamate conjugated to bovine serum albumin (BSA), KLH and shows no cross-reactivity with L-aspartate, L-glutamine, L-asparagine, L-alanine or BSA, and only weak cross-reactivity with glycyl-L-aspartic acid, GABA, β-alanine, glycine and 5-aminovaleric acid. Pre-incubation of spinal cord slices with glutamate eliminates G6642 immunoreactivity in zebrafish [9]. Immunostaining with G6642 labels the majority of neurons expressing a fluorescent protein under control of the Vglut2 promoter [9] or under control of the Chx10 promoter in zebrafish (i.e. V2a neurons, a population of spinal glutamatergic interneurons that generate the locomotor rhythm from fish to mammals, [10]). Double labeling with Neurotrace fluorescent Nissl stain indicated that cells labelled by G6642 are neurons in zebrafish spinal cord [10]. The G6642 antibody was successfully used to stain glutamate in photoreceptor cells in mouse retina [8], glutamatergic neurons in rat brainstem [11, 12] or to distinguish glutamatergic from glycinergic and/or GABAergic populations in zebrafish spinal cord [10], rat auditory brainstem circuits [13] and mouse amygdala [14].

The AB144P ChAT antibody has been widely used to label cholinergic neurons in lamprey [15, 16, 17], salamander [18, 19, 20], rat [21], human [22, 21] and mouse brain tissues [23]. This affinity purified polyclonal antibody is raised against the human placental enzyme. The supplier has tested its specificity in human placenta lysates and using western blots on mouse brain lysates, where it detects a band of 68-70 kDA. It labels neurons expressing a fluorescent protein under control of the ChAT promoter in mice [24].

### Specificity of the transgenic mice

#### Vglut2-ires-Cre mouse

These mice are widely used to express Cre-recombinase in glutamatergic Vglut2-positive neurons without interfering with *Vglut2* gene expression [25]. When Vglut2-ires-Cre are crossed with a lox-GFP mouse, GFP-positive neurons are found in glutamatergic regions (positive for *Vglut2* mRNA) and absent from GABAergic regions (positive for the vesicular GABA transporter mRNA) [25]. When Vglut2-ires-Cre are crossed with a lox-tdTomato mouse, the cells labelled in the dorsal horn of the spinal cord are immuno-positive for NeuN and immuno-negative for Pax2 and for Wilm’s tumor 1, two markers of inhibitory neurons [26, 27]. Chemogenetic activation of Vglut2-Cre neurons increases the frequency of synaptic excitatory currents recorded with patch-clamp in spinal cord slices [26] and evokes short latency excitatory responses in periaqueductal gray neurons [28]. Excitatory postsynaptic responses are evoked in the striatum when stimulating thalamic terminals in mice obtained by crossing the Vglut2-ires-Cre with lox-channelrhodopsin (**ChR2**) mice [29]. The Vglut2-ire-Cre mouse was used to study the role of reticulospinal neurons in locomotor control [30].

#### ZsGreen-lox mouse

The Ai6 mouse has been widely used to label cells expressing the Cre-recombinase [31]. After exposure to Cre-recombinase, the floxed STOP cassette is removed, and this results in the expression of the ZsGreen fluorescent protein under control of CAG promoter. Cells display intense labelling with ZsGreen as demonstrated by us [1] and others [32, 23]. In our previous study, we compared ZsGreen-lox mouse (Cre-negative) and Vglut2-ZsGreen (Cre-positive) brain sections and confirmed that prior to introduction of Cre-recombinase, only a very low baseline level of fluorescence was present in brain slices of homozygous ZsGreen-lox mice [1]. This is classical for reporter lines based on CAG promoter-driven expression (e.g. Ai9, tdTomato-lox mouse) as mentioned by the supplier (Jackson laboratory).

#### ChR2-EYFP-lox mouse

The Ai32 mouse [31] has been widely used to activate cells expressing the Cre-recombinase using optogenetics (e.g. [33, 34]). When exposed to Cre-recombinase, the floxed STOP cassette is removed, and this results in the expression of the ChR2(H134R)-EYFP fusion protein under control of CAG promoter.

### DeepLabCut networks

For open-field locomotion analysis, we labelled 6 landmarks on 520 frames taken from 20 videos of 8 different animals assigning the 95% of those images to the training set without cropping. The landmarks were the body center, the four corners of the arena, and the low-power LED to visualize optogenetic stimulation. We used a ResNet-50-based neural network [35, 36] with default parameters for 1,030,000 training iterations. We validated with one shuffle and found the test error was 2.28 pixels and the train error 1.85 pixels.

For footfall pattern analysis, we labelled 9 landmarks 489 frames taken from 27 videos of 9 animals assigning the 95% of those images to the training set without cropping. The landmarks were the four paws, the four distance calibration markers, and the low-power LED to visualize optogenetic stimulation. We used a ResNet-50-based neural network [35, 36] with default parameters for 1,030,000 training iterations. We validated with one shuffle and found the test error was 2.31 pixels and the train error 1.76 pixels.

For limb kinematics analysis, we labelled 11 landmarks on 906 frames taken from 44 videos of 7 different animals assigning the 95% of those images to the training set without cropping. The landmarks were the 5 joints and the toe tip, the four distance calibration markers, and the low-power LED to visualize optogenetic stimulation. We used a ResNet-50-based neural network [35, 36] with default parameters for 1,030,000 training iterations and one refinement of 1,030,000 iterations. We validated with one shuffle and found the test error was 2.03 pixels and the train error 1.87 pixels.

